# The Deubiquitylating Enzyme Ubp12 Regulates Rad23-Dependent Substrate Delivery at the Proteasome

**DOI:** 10.1101/108068

**Authors:** Daniela Gödderz, Nico P. Dantuma

## Abstract

The consecutive actions of the ubiquitin-selective segregase Cdc48 and the ubiquitin shuttle factor Rad23 result in the delivery of ubiquitylated proteins at the proteasome. Here, we show that the deubiquitylating enzyme Ubp12 interacts with Cdc48 and regulates proteasomal degradation of Rad23-dependent substrates. Overexpression of Ubp12 results in stabilization of Rad23-dependent substrates. We show that Ubp12 removes short ubiquitin chains from the N-terminal ubiquitin-like domain (UbL) of Rad23. Preventing ubiquitylation of Rad23 by substitution of lysine residues within the UbL domain, Rad23^UbLK0^, does not affect the non-proteolytic role of Rad23 in DNA repair but causes an increase in ubiquitylated cargo bound to the UBA2 domains of Rad23 and recapitulates the stabilization of Rad23-dependent substrates observed upon overexpression of Ubp12. Expression of Rad23^UbLK0^ or overexpression of Ubp12 impairs the ability of yeast to cope with proteotoxic stress consistent with inefficient clearance of misfolded proteins by the ubiquitin/proteasome system. Our data suggest that ubiquitylation of Rad23 plays a stimulatory role in facilitating the transfer of ubiquitylated substrates to the proteasome.

**Summary statement:** Ubiquitylation of a ubiquitin shuttle factor regulates the delivery of substrates at the proteasome, uncovering a novel regulatory link between ubiquitin and proteasomal degradation.

## Introduction

Posttranslational modifications with the protein modifier ubiquitin are best known for their critical role in targeting proteins for proteasomal degradation (Hershko and Ciechanover, 1998). More recent work has revealed the existence of an elaborated ubiquitin code that orchestrates not only proteasomal degradation but also a broad array of other cellular processes by regulating protein-protein interactions through ubiquitylation (Komander and Rape, 2012). Because of the wealth of knowledge on the ubiquitin/proteasome system (UPS) and its many players, this essential proteolytic system remains an important paradigm for understanding the spatial and temporal control of interactions between ubiquitylated proteins and ubiquitin binding proteins. Although ubiquitin modifications have been originally considered as a point of no return, leading inevitably to the destruction of the tagged protein, it has become apparent that a number of proteins not only handle ubiquitylated proteins but can also change their fate (Crosas et al., 2006, Hanna et al., 2006, Richly et al., 2005, Verma et al., 2004). These proteins that share the ability to interact with ubiquitylated proteins – either through the presence of ubiquitin-binding domains or by interacting with proteins that contain these motifs – can depending on their mode of action either promote or prevent the degradation of ubiquitylated proteins.

In yeast, the ubiquitin-selective segregase Cdc48 (also known as valosin-containing protein (VCP) or p97 in mammals) is an interaction hub for proteins that modulate ubiquitylated proteins (Jentsch and Rumpf, 2007). While the intrinsic segregase activity of this AAA-ATPase appears to prepare proteins for destruction by extracting them from macromolecular complexes (Dai et al., 1998, Elkabetz et al., 2004) and/or generating unfolded structures that allow efficient proteasomal degradation (Beskow et al., 2009), it also associates with a number of proteins, including ubiquitin ligases, ubiquitin elongation factors and deubiquitylating enzymes (DUBs), that modify the ubiquitin signal on the substrates (Richly et al., 2005). Moreover, Cdc48 binds the ubiquitin shuttle factor Rad23 (Baek et al., 2011), which sequesters ubiquitylated proteins and delivers them at the proteasome (Schauber et al., 1998). Critical for the interaction of Rad23 with Cdc48 and the proteasome is Rad23’s ubiquitin-like domain (UbL) domain, which resembles ubiquitin but has distinct properties (Lambertson et al., 2003). It has been proposed that in a sequential mode Cdc48 and Rad23 facilitate the destruction of proteasome substrates by ensuring that they reach the proteasome in a state allowing efficient degradation (Richly et al., 2005). Consistent with this model, insertion of a short peptide extension that is sufficiently long to function as an unstructured initiation site abrogates the need for both Cdc48 and Rad23 in proteasomal degradation (Godderz et al., 2015).

Rad23 interacts with the proteasome by means of its N-terminal UbL domain while the ubiquitin-associated (UBA)-2 domain located at its most extreme C-terminus functions as an intrinsic stabilization signal (Heessen et al., 2005) that hinders initial unfolding events at the entrance of the proteasome, allowing Rad23 to resist proteasomal degradation (Fishbain et al., 2011, Heinen et al., 2011). This endows Rad23 to function as a reusable shuttle that delivers its ubiquitylated cargo, which is bound to the ubiquitin binding UBA1 and UBA2 domains, at the proteasome where the cargo will selectively bind to the ubiquitin receptors that reside in the regulatory particle of the proteasome (Finley, 2009). Rad23 shares its molecular structure and function with Dsk2 and Ddi1, two other ubiquitin shuttle factors with distinct but overlapping substrate specificities (Crosas et al., 2006). These three ubiquitin shuttle factors potentially provide an additional layer of specificity in substrate targeting which may enable the cell to prioritize a specific set of substrates.

Although it is presently unclear if the functionality of ubiquitin shuttle factors is differentially regulated, there are at least two molecular mechanisms that may control the activity of these scaffold proteins. Firstly, specific binding proteins may interact with the functional domains in the ubiquitin shuttle proteins and prevent them from collecting and/or delivering their ubiquitylated cargo. In this respect, it is noteworthy that the peptidyl-tRNA hydrolase Pth2 has been reported to interact with the UbL domain of Rad23 and Dsk2 restricting their access to the proteasome (Ishii et al., 2006). Moreover, the extraproteasomal population of the ubiquitin-binding receptor Rpn10 selectively binds Dsk2 and prevents it from docking to the proteasome (Matiuhin et al., 2008). Secondly, posttranslational modifications may change the behavior of the ubiquitin shuttle factors and may provide a means to control their activity. Notably, phosphorylation of the UbL domain of Rad23 regulates its interaction with the proteasome (Liang et al., 2014), while modification of Dsk2 with proteolytic ubiquitin chains results in a reduced capacity to bind polyubiquitylated proteins (Sekiguchi et al., 2011). Moreover, recruitment of the ubiquitin-binding receptor Rpn10 is regulated by monoubiquitylation (Isasa et al., 2010). Thus, the functional status of ubiquitin shuttle factors can be modulated by protein-protein interactions and posttranslational modifications.

In the present study, we show that Rad23 is modified in its UbL domain with short polyubiquitin chains and identified Ubp12 as a Cdc48-interacting DUB that selectively reverses this modification. Interestingly, abrogating Rad23 ubiquitylation causes an increase in ubiquitylated cargo associated with Rad23, which, despite being proficient for binding to the proteasome, fails to facilitate proteasomal degradation of the substrates. Our data show that ubiquitylation of Rad23 is important for proper proteasomal degradation and suggest that this Ubp12-controlled mechanism may control the release of ubiquitylated proteins at the proteasome.

## Results

### The DUB Ubp12 interacts with Cdc48 and stabilizes Rad23-dependent substrates

Yeast Cdc48 and the mammalian orthologue VCP/p97 are known to interact with a number of proteins involved in ubiquitylation, including the deubiquitylating enzymes Otu1 (Rumpf and Jentsch, 2006) and ataxin-3 (Wang et al., 2006), respectively. We found that also the poorly characterized ubiquitin-specific protease Ubp12 interacts with Cdc48 in the budding yeast *Saccharomyces cerevisiae*. We observed that immunoprecipitation of endogenous Cdc48 resulted in co-purification of HA-tagged Ubp12 (Ubp12^6HA^) expressed from the endogenous promoter (Fig. 1A). This interaction was further validated by showing in the reciprocal immunoprecipitation that endogenous Cdc48 was pulled down with FLAG-tagged Ubp12 (^FLAG^Ubp12) (Fig. 1B).

**Figure 1.**
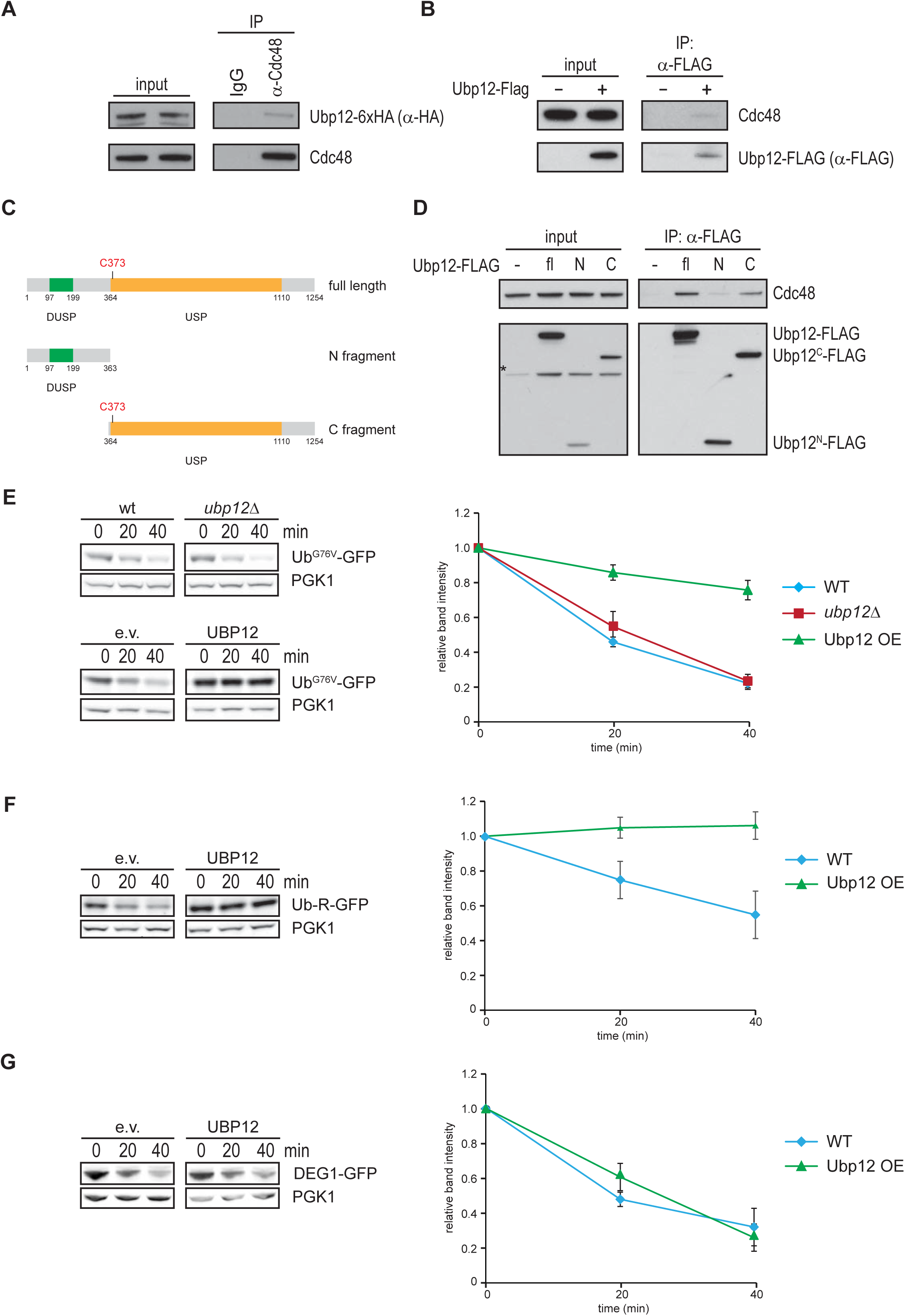
The DUB Ubp12 interacts with Cdc48 and stabilizes Rad23-dependent substrates. **A)** Endogenous Cdc48 was immunoprecipitated (IP) with a Cdc48-specific antibody from wild-type yeast expressing endogenously 6xHA-tagged Ubp12. Non-specific IgG was used as control. Immunoprecipitated proteins were analyzed by immunoblotting using antibodies indicated on the left. As a reference 2% of total input lysate was analyzed in parallel. **B)** Immunoprecipitation was performed with an anti-FLAG antibody using lysates from a *ubp12*Δ yeast that did or did not overexpress Ubp12-Flag. Non-specific IgG was used as control. Immunoprecipiated proteins were probed with the indicated antibodies. As a reference 2% of total input lysate was analyzed in parallel. **C)** Schematic representation of the domain structure of the Ubp12 protein. Two fragments were generated for characterization of Ubp12-Cdc48 interaction. **D)** Anti-FLAG IP was performed as in B from a *ubp12*Δ yeast overexpressing either FLAG-tagged full-length Ubp12, the N- or the C-terminal fragment as depicted in *C*. Input controls correspond to 2% of total lysate. Immunoprecipitated proteins were analyzed by immunoblotting using antibodies indicated on the left. A cross-reactive band is marked with an asterisk. **E)** Turnover of the UFD model substrate Ub^G76V^-GFP was determined by cycloheximide chase in wild-type yeast containing either empty vector (e.v.) or overexpressing Ubp12 from a high-copy plasmid or *ubp12*Δ yeast. Proteins were analyzed by immunoblotting with a GFP- and PGK1-specific antibodies. Quantification of GFP levels normalized to PGK1 of three independent experiments with error bars representing standard error of the mean is shown. **F)** As in (E), turnover of N-end rule model substrate Ub-R-GFP was analyzed in wild-type yeast that did or did not overexpress Ubp12-FLAG. **G)** As in (E), turnover of DEG1-GFP was analyzed in wild-type yeast that did or did not overexpress Ubp12-FLAG.

Ubp12 is a member of the ubiquitin-specific protease (Ubp) family that is comprised of 16 members in yeast, none of which are essential for viability (Reyes-Turcu et al., 2009). These proteins share a conserved catalytic core domain and possess terminal extensions and insertions. Ubp12 is a large protein containing in its N-terminal part a DUSP domain, suggested to be important for protein-protein interactions (Song et al., 2010), while the USP catalytic domain comprises the central part of the protein (Fig. 1C). To further characterize the interaction of Ubp12 with Cdc48, two constructs expressing FLAG-tagged fragments of Ubp12 either containing the N-terminal DUSP domain (Ubp12-N^FLAG^) or the C-terminal USP domain (Ubp12-C^FLAG^) were generated. Interestingly, Cdc48 primarily interacted with the C-terminal fragment that included the catalytic USP domain (Fig. 1D).

To understand the functional importance of the Ubp12-Cdc48 interaction, we analyzed the degradation of Cdc48-dependent proteasomal substrates. Cdc48 has originally been implicated in the ubiquitin/proteasome system by its identification as a critical factor for proteasomal degradation of ubiquitin fusion degradation (UFD) substrates (Johnson et al., 1995) and N-end rule substrates (Ghislain et al., 1996). In line with a role of Ubp12 in degradation of Cdc48-dependent substrates, turnover analysis revealed that the UFD substrate Ub^G76V^-GFP was stabilized upon overexpression of Ubp12 (Fig. 1E), while expression of the catalytically inactive Ubp12^C372S^ mutant did not delay degradation indicating that its deubiquitylating activity is required for the stabilizing effect (**Suppl. Fig. S1**). Also degradation of the N-end rule substrates Ub-R-GFP, which is targeted by its N-terminal arginine residue upon cleavage of the ubiquitin moiety by ubiquitin C-terminal hydrolyses (Varshavsky, 1996), was impaired upon overexpression of Ubp12 (Fig. 1F). Surprisingly, the Cdc48-dependent model substrate DEG1-GFP, which is based on the natural degradation signal of the yeast mating factor Matα2 (Chen et al., 1993), was not affected by Ubp12 overexpression (Fig. 1G). Together these data show that Ubp12 is a Cdc48-interacting DUB that stabilizes some, but not all, Cdc48-dependent substrates.

### Ubp12 modulates Rad23 ubiquitylation

We noticed that a difference between these Cdc48-dependent substrates is their reliance on the ubiquitin shuttle factor Rad23 for proteasomal degradation. Thus, whereas Ub^G76V^-GFP (**Suppl. Fig. S2A**) and Ub-R-GFP (**Suppl. Fig. S2B**), which are stabilized by Ubp12 overexpression, require Rad23 for degradation (Godderz et al., 2015), DEG1 is degraded in a Rad23-independent fashion (Medicherla et al., 2004). Since Rad23 operates downstream of Cdc48 by delivering ubiquitylated substrates at the proteasome (Richly et al., 2005), an effect of Ubp12 on Rad23-dependent degradation would be consistent with a stabilizing role of Ubp12 in a specific branch of the Cdc48 pathway. To test a possible function of Ubp12 in Rad23-dependent degradation, we determined if Ubp12 physically interacts with Rad23. Indeed, we found that immunoprecipitation of Ubp12^FLAG^ resulted in the co-precipitation of Rad23 (Fig. 2A). This interaction was further corroborated by the reciprocal immunoprecipitation in which Ubp12^FLAG^ was pulled down with endogenous Rad23 (Fig. 2B). Interestingly, the interaction with Ubp12 was strictly dependent on the N-terminal UbL domain of Rad23 (Fig. 2C).

**Figure 2.**
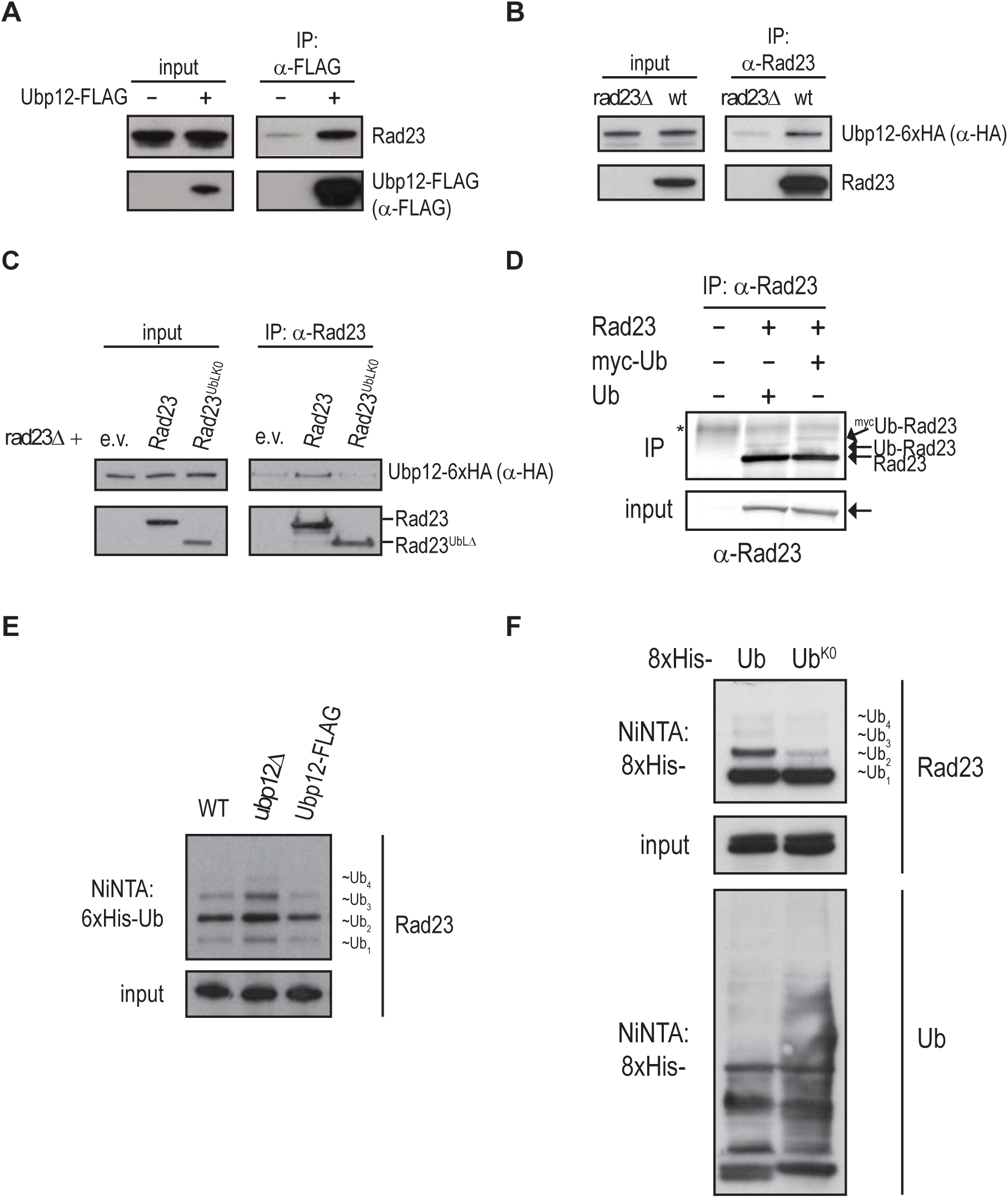
Ubp12 modulates Rad23 ubiquitylation. **A)** Anti-FLAG immunoprecipitation (IP) was performed from a *ubp12*Δ yeast that did or did not overexpress Ubp12-FLAG. Input controls correspond to 2% of total lysate. Immunoprecipitated proteins were analyzed by immunoblotting using antibodies indicated on the left. **B)** Immunoprecipitation with a Rad23-specific antibody was performed from wild-type or *rad23*Δ yeast expressing endogenously 6xHA-tagged Ubp12. Immunoprecipitated proteins were analyzed by immunoblotting using antibodies indicated on the left. Input controls correspond to 2% of total lysate. ***C)*** Anti-Rad23 immunoprecipitation was performed as in (B) from *rad23Δ* yeast with endogenously 6xHA-tagged Ubp12 and containing low-copy plasmids expressing either full-length Rad23 or Rad23 lacking the UbL domain (Rad23^UbLΔ^). Input controls correspond to 2% of total lysate. Immunoprecitated proteins were analyzed by immunoblotting using antibodies indicated on the left. **D)** Anti-Rad23 IP was performed from either wild-type yeast or yeast expressing myc-tagged ubiquitin (myc-Ub) as the sole source for ubiquitin. Immunoprecipitated proteins were analyzed by immunoblotting using Rad23-specific antibody. **E)** Denaturing Ni-NTA pulldown from wild-type, *ubp12*Δ or Ubp12-FLAG overexpressing yeast co-expressing 6His-tagged ubiquitin (6His-Ub) was performed. Immunoprecipitated proteins were analyzed by immunoblotting using a Rad23-specific antibody. Input controls correspond to 2% of total lysate. **F)** Denaturing Ni-NTA pulldown from wild-type yeast either expressing 8xHis tagged wild-type (Ub) or lysine less ubiquitin (Ub^K0^) was performed. Precipitated proteins were analyzed by immunoblotting using antibodies indicated on the right.

Rad23 has been found in a large-scale mass spectrometric analysis to be ubiquitylated at multiple lysine residues in its UbL domain but the functional significance of these modifications has remained elusive (Swaney et al., 2013). Immunoprecipitation of endogenous Rad23 indeed revealed the presence of a band that corresponded in size to ubiquitylated Rad23 and that was shifted to a slightly higher molecular weight in yeast overexpressing Myc-tagged ubiquitin (^Myc^Ub), consistent with these bands representing Rad23 modified with wild-type or epitope-tagged ubiquitin, respectively (Fig. 2D). To validate the presence of ubiquitylated Rad23 in yeast lysates, we precipitated His-tagged ubiquitin (^His^Ub) under denaturing conditions using NiNTA beads in order to avoid co-immunoprecipitation of non-covalently bound ubiquitin conjugates and probed the precipitated ubiquitin conjugates for endogenous Rad23. This more sensitive method revealed Rad23 species that were modified with up to four ubiquitin molecules, confirming that Rad23 is covalently modified with multiple ubiquitin moieties (Fig. 2E). We wondered if the interaction between Ubp12 and Rad23 had an effect on the ubiquitylation status of Rad23. Indeed, we found that the steady-state levels of these ubiquitin-modified Rad23 species were increased in a strain lacking Ubp12 (*ubp12Δ*) and decreased upon overexpression of Ubp12 in line with Ubp12 being a DUB that targets Rad23 ubiquitylation (Fig. 2E). In order to investigate whether this activity is unique to Ubp12, we analyzed the ubiquitylation status of Rad23 using deletion strain for each of the deubiquitylating enzymes belonging to the Ubp family with the exception of Ubp8 (data not shown). We found that only deletion of Ubp3 also resulted in an increase of ubiquitylated Rad23 (**Suppl. Fig. S3A**), but concluded that this was likely to be an indirect effect since this was, unlike the situation in the *ubp12Δ* strain, accompanied by a general accumulation of ubiquitin conjugates (**Suppl. Fig. S3B**). Moreover, simultaneous deletion of Ubp3 and Ubp12 resulted in a further increase suggesting that Ubp3 causes an increase in ubiquitylated Rad23 by a different mechanism than Ubp12 consistent with the general effect on ubiquitin conjugates (**Suppl. Fig. S3A**). Thus, the ability to selectively reverse ubiquitylation of Rad23 appears to be a unique property of Ubp12.

Instead of a smear of high molecular weight ubiquitylated species, which is typically observed for polyubiquitylated substrates, we predominantly detected a limited number of distinct bands that corresponded in size with Rad23 being modified with one to four ubiquitin moieties. To discriminate whether these ubiquitin-modified species represented Rad23 modified with a short ubiquitin chain or Rad23 that was monoubiquitylated at multiple lysine residues, we analyzed whether overexpression of a His-tagged lysine-less ubiquitin (^His^Ub^K0^) affected the abundance of these modified species. If Rad23 is subject to multiple monoubiquitylation the expression of ^His^Ub^K0^ should not affect their relative abundance. However, in the case of short ubiquitin chains the band corresponding to a single ubiquitin conjugated to Rad23 should not be affected while the larger species are expected to be reduced since the ^His^Ub^K0^ will function as a chain terminator. In line with the latter possibility, the bands corresponding to Rad23 with two, three and four ubiquitin moieties were reduced in the presence of ^His^Ub^K0^ while the monoubiquitylated Rad23 remained at the same level (Fig. 2F). Together our data indicate that Rad23 is modified with short ubiquitin chains that are negatively regulated by Ubp12.

### The UbL domain of Rad23 is ubiquitylated

We aimed to identify the lysine residue within Rad23 that is modified with the ubiquitin chain. Rad23 contains fifteen lysine residues of which all, except for one, are located in the N-terminal UbL domain (Fig. 3A). Supporting the idea that the UbL domain is the target for this ubiquitylation event, we observed that deletion of this domain abrogated ubiquitylation of Rad23 (Fig. 3B). In order to identify the lysine residue(s) that serves as ubiquitin acceptor site(s) in Rad23, we generated a collection of Rad23 mutants in which one or several lysine residues were simultaneously replaced by arginine residues (**Suppl. Fig. S4**). Analysis of the ubiquitylation status of these Rad23 variants showed that all mutants containing lysine residues in the UbL domain were subject to ubiquitin conjugation while ubiquitylation did not occur when all fourteen lysine residues in the UbL domain had been replaced by arginine residues (Rad23^UbLK0^) (Fig. 3C). This suggested that the short ubiquitin chain that modifies Rad23 can be conjugated in a promiscuous fashion to multiple lysine residues within the UbL domain suggesting a relaxed stringency of the ubiquitin ligase(s) involved. To empirically address this, we compared the ubiquitylation pattern of three Rad23 mutants that each contained a single lysine residue at different positions in the UbL domain. Each of these Rad23 mutants was ubiquitylated confirming that a single lysine residue in various positions can restore ubiquitylation of Rad23 (Fig. 3D).

**Figure 3.**
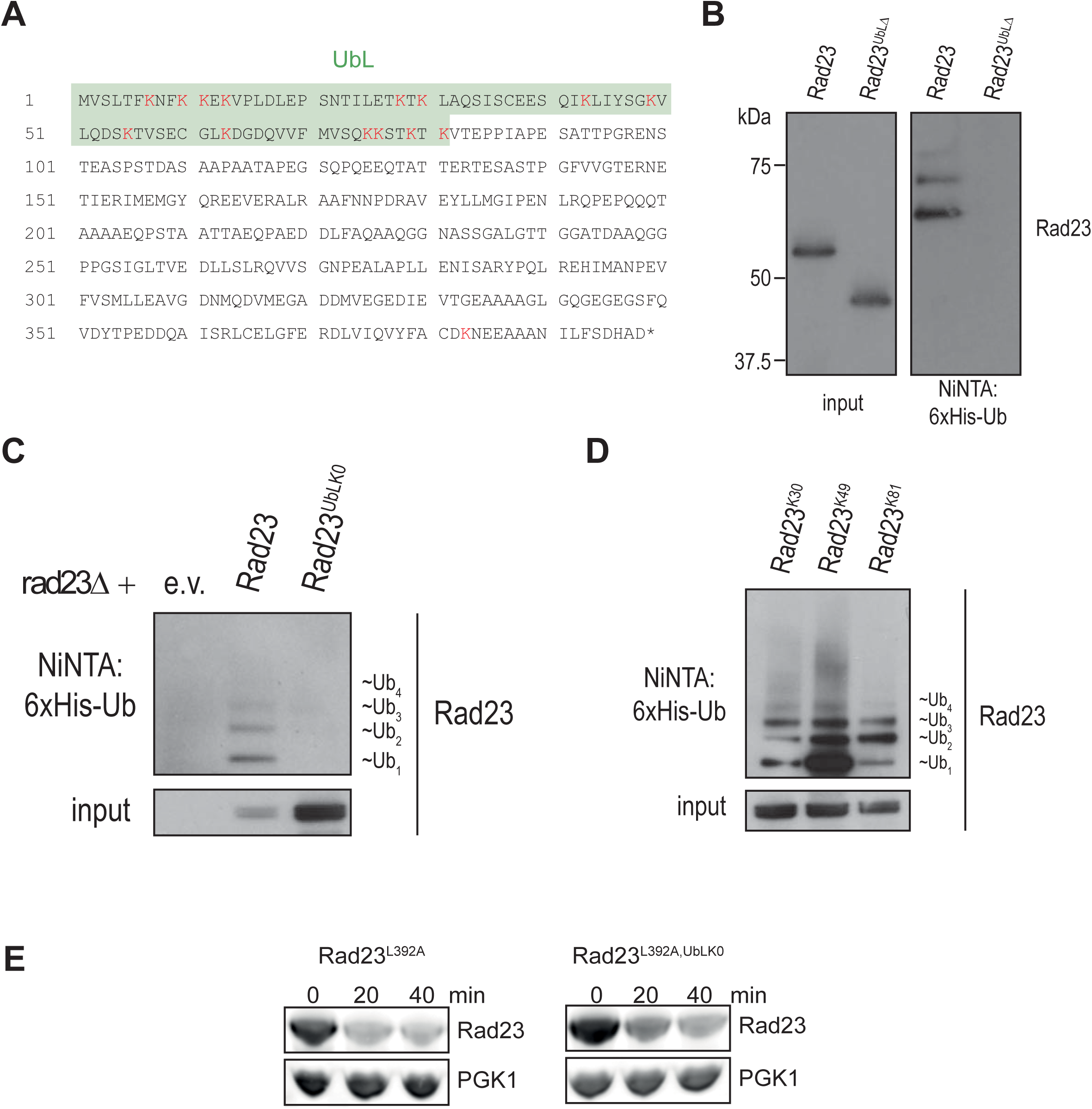
The UbL domain of Rad23 is ubiquitylated. **A)** Amino acid sequence of Rad23 is depicted with UbL domain marked by green shading and lysine residues highlighted in red. **B)** Denaturing Ni-NTA pulldown from *rad23*Δ yeast co-expressing 6His-Ub and either full-length Rad23 or Rad23^UbLΔ^ was performed. Precipitated proteins were analyzed by immunoblotting using Rad23-specific antibody. **C)** Denaturing Ni-NTA pulldown from *rad23*Δ yeast cells co-expressing 6His-Ub and either wild-type Rad23 or a variant with all lysine residues in the UbL domain mutated to arginine (Rad23^UbLK0^) was performed. Bound proteins were analyzed by immunoblotting using Rad23-specific antibody. **D)** As in (B) but yeast expressed Rad23 variants in which single lysine residues as indicated had been reintroduced. **E)** Turnover of Rad23^L392A^ and Rad^L392A,UbLK0^ was determined by cycloheximide chase in wild-type yeast. The membranes were probed with Rad23- and PGK1-specific antibodies.

The UbL domain of Rad23 can function analogous to a degradation signal without resulting in actual degradation of Rad23 because of its C-terminal UBA2 domain, which acts as an intrinsic stabilization degradation that prevents degradation (Heessen et al., 2005), by hindering the initiation of unfolding events required for proteasomal degradation (Fishbain et al., 2011, Heinen et al., 2011). We wondered if these ubiquitin chains could be responsible for the ability of the UbL domains to target Rad23 for proteasomal degradation upon inactivation of the protective UBA2 domain. However, substitution of all the lysine residues in the UbL domain, in contrast to deletion of the UbL domain, did not result in stabilization of Rad23^L392A^, which lacks the protective UBA2 domain, suggesting that these ubiquitin chains do not contribute to targeting Rad23^L392A^ for proteasomal degradation (Fig. 3E). This also shows that the mutant UbL domain lacking all lysine residues is proficient in targeting Rad23 for degradation and therefore still capable to interact with the proteasome.

### Preventing Rad23 ubiquitylation stabilizes substrates and enhances levels of ubiquitylated cargo

To investigate the role of ubiquitylation in proteasomal degradation, we compared the functionality of Rad23 and Rad23^UbLK0^ in the turnover of the Rad23-dependent substrate Ub^G76V^-GFP and Rad23-independent substrate DEG1-GFP. This revealed that wild-type Rad23 fully restored degradation of Ub^G76V^-GFP in a *rad23Δ* strain while Rad23^UbLK0^ only had a partial rescuing effect that was comparable in magnitude to what was observed upon Ubp12 overexpression (Fig. 4A). As expected deletion of Rad23 did not affect the degradation of DEG1-GFP but, notably, degradation was neither distorted by overexpression of Rad23^UbLK0^ again suggesting that this mutant does not cause a general impairment of ubiquitin-dependent degradation (Fig. 4B). Thus, substitution of lysine residues in the UbL domain impairs the ability of Rad23 to facilitate degradation of specific substrates, without causing a general impairment of proteasomal degradation, in line with a function of Rad23 ubiquitylation in proteasomal degradation.

**Figure 4.**
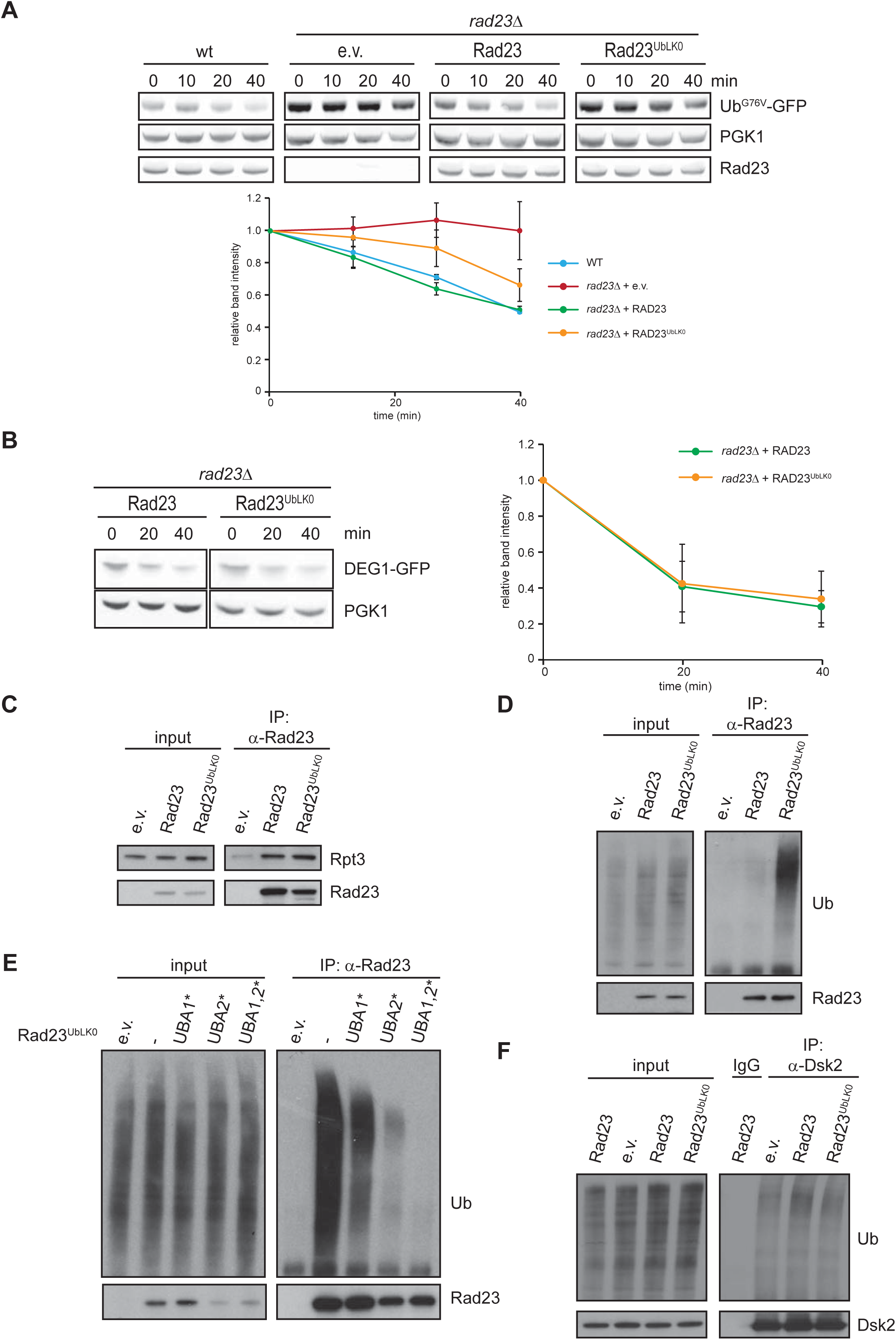
Preventing Rad23 ubiquitylation stabilizes substrates and enhances levels of ubiquitylated cargo. **A)** Turnover of Ub^G76V^-GFP was determined by cycloheximide chase in wild-type or *rad23Δ* yeast cells containing either empty vector (e.v.) or a low-copy plasmid expressing Rad23 or Rad23^UbLK0^. Proteins were analyzed by immunoblotting. Quantification of GFP levels normalized to PGK1 of three independent experiments with error bars representing standard error of the mean is shown. **B)** Turnover of DEG1-GFP was monitored by cycloheximide chase in *rad23Δ* yeast transformed with either an expression vector for Rad23 or Rad23^UbLK0^. Quantification of GFP levels normalized to PGK1 of three independent experiments with error bars representing standard error of the mean is shown. **C)** Rad23 was immunopreciptated from *rad23Δ* yeast expressing plasmid-derived wild-type Rad23 or Rad23^UbLK0^. Immunopecipiated proteins were probed for the proteasome subunit Rpt3. The input control corresponds to 2% of total lysate. **D)** To analyze binding of ubiquitylated cargo to Rad23, Rad23 was immunoprecipitated from *rad23Δ* strain containing low-copy plasmids expressing the indicated Rad23 variants. Immunoprecipiated proteins were analyzed by immunoblotting using antibodies against Rad23 and ubiquitin. The input control corresponds to 2% of total lysate. **E)** As in (D) but here Rad23 variants in which the UBA1, UBA2 or UBA1 and UBA2 had been mutated to abrogate their ubiquitin binding properties. **F)** Dsk2 was immunoprecipitated from a *rad23Δ* strain containing low-copy plasmids expressing the indicated Rad23 variants. IgG was used for control IP. Bound proteins were analyzed by immunoblotting using antibodies against Dsk2 and ubiquitin. The input control corresponds to 2% of total lysate.

Since Rad23 functions as a scaffold protein that mediates degradation by simultaneously binding the proteasome and ubiquitylated substrates, we argued that the stabilizing effect of Rad23 ubiquitylation was likely due to either an impaired binding to the proteasome or an altered interaction with ubiquitylated substrates. The above-mentioned observation that substitution of lysine residues in the UbL domain does not stabilize Rad23^L392A^ argues against modulation of the Rad23-proteasome interaction. To more directly address this question, we immunoprecipitated Rad23 and Rad23^UbLK0^ from yeast lysates, which indeed confirmed that both interacted with the proteasome (Fig. 4C). This suggests that ubiquitylation is not stimulating or hindering the binding of the UbL domain to the proteasome and at the same time confirms that the overall structural integrity of the UbL domain is not affected by the lysine substitutions.

We investigated if binding of ubiquitylated cargo to Rad23 was altered in the absence of lysine residues in the UbL domain. Interestingly, we observed a strong increase in ubiquitylated cargo bound to Rad23^UbLK0^ (Fig. 4D). Introduction of mutations in the two UBA domains of Rad23, which are responsible for the binding of ubiquitylated proteins (Bertolaet et al., 2001), almost completely abolished co-immunoprecipitation of ubiquitin conjugates suggesting that the UbLK0 domain increases the load of ubiquitylated proteins bound to the UBA domains of Rad23 (Fig. 4E). A possible explanation could be that Rad23^UbLK0^ impairs the degradation of ubiquitylated proteins resulting in a global increase in ubiquitin conjugates which may, in turn, increase the amount of substrates that are captured by Rad23. However, we did not observe a general increase in ubiquitin conjugates in yeast expressing Rad23^UbLK0^ arguing against this scenario (Fig. 4E). Moreover, if this would be the case, also other ubiquitin shuttle factors are expected to display an increase in the load of polyubiquitylated substrates. To probe into this, we immunoprecipitated the related ubiquitin shuttle factor Dsk2 in yeast expressing either wild-type Rad23 or Rad23^UbLK0^. Unlike the situation for Rad23, we found that the levels of ubiquitylated proteins that were bound to Dsk2 were comparable under both conditions (Fig. 4F). We conclude therefore that preventing ubiquitylation of the UbL domain of Rad23 does not interfere with the UbL-dependent binding of Rad23 to the proteasome but selectively increases the pool of ubiquitylated substrates that is associated with the UBA domains of Rad23.

### Rad23^UbLK0^ is proficient in DNA repair but defective in proteasomal degradation

Rad23 does not only play important roles in proteasomal degradation but is also involved in nucleotide excision repair, which is the primary pathway for removing helix-distorting lesions from DNA (Dantuma et al., 2009). Although the latter is dependent on the UbL domain of Rad23 and involves also the proteasome, it is believed to be a largely non-proteolytic process as it does not require the degradation capacity of the proteasome (Russell et al., 1999). We observed that the ultraviolet light (UV) sensitivity of the *rad23Δ* strain could be rescued by expression of wild-type Rad23 but not by Rad23 lacking the UbL domain (Rad23^UbLΔ^) consistent with a critical role of the UbL domain in nucleotide excision repair (Fig. 5A). However, expression of Rad23^UbLK0^ gave a comparable rescue of the viability of yeast upon UV exposure to wild-type Rad23 demonstrating that the UbLK0 domain is functional in DNA repair and arguing against a role for Rad23 ubiquitylation in nucleotide excision repair.

**Figure 5.**
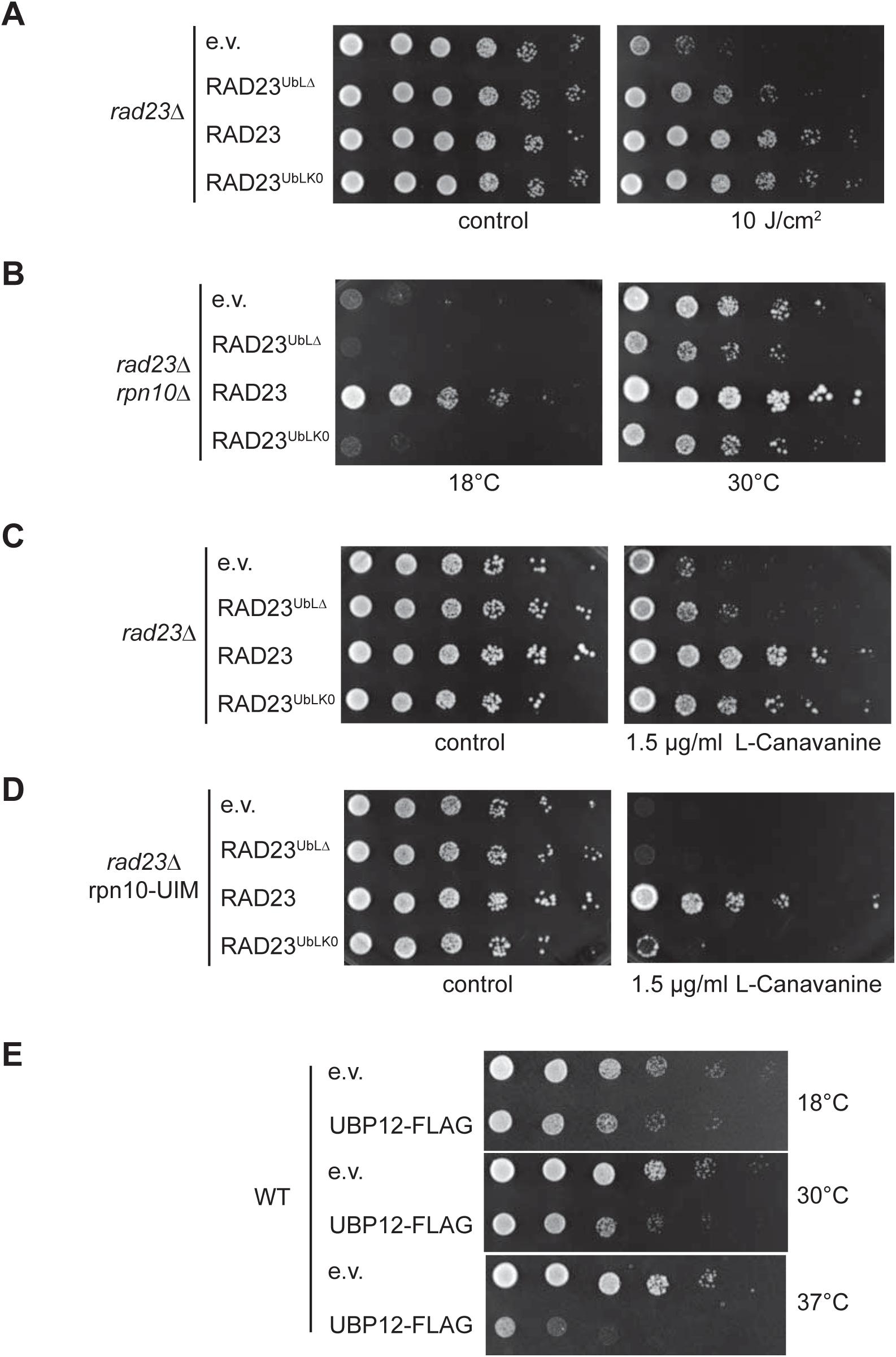
Rad23^UbLK0^ is proficient in DNA repair but defective in proteasomal degradation. **A**) The *rad23Δ* yeast was transformed with empty vector (e.v.) or plasmids expressing Rad23^UbLΔ^, Rad23 or Rad23^UbLK0^. Yeast cultures were grown to exponential phase, and 6-fold dilutions were spotted on synthetic medium and exposed to UV light of the indicated intensity prior to incubation at 30ºC for three days. **B)** The *rad23Δ/rpn10Δ* strain was transformed with empty vector (e.v.) or plasmids expressing Rad23^UbLΔ^, Rad23 or Rad23^UbLK0^. Yeast cultures were grown to exponential phase, and 6-fold dilutions were spotted on synthetic medium and grown at 18°C or 30°C. Growth was examined after 3 to 5 days of incubation, depending on the temperature. C) As in (A) but yeast was spotted on synthetic medium or synthetic medium containing 1.5 μg/ml L-canavanine and incubated at indicated temperatures. **D)** As in (C) but *rad23Δ*/rpn10-UIM yeast was used. **E)** Wild-type yeast was transformed with an empty vector (e.v.) or an expression vector for Ubp12-FLAG. Growth was examined at 18°C, 30°C and 37°C.

Due to redundancy of the ubiquitin shuttle factors and the intrinsic ubiquitin receptors in the proteasome in substrate delivery, deficiencies in any of these factors give relatively mild phenotypes unless multiple delivery proteins are simultaneously compromised. To unmask effects of Rad23^UbLK0^ on proteasomal degradation, we studied therefore the effect of this Rad23 variant in yeast that lacked the ubiquitin receptor Rpn10 proteasome subunit. Deletion of both Rpn10 and Rad23 results in a mild growth impairment that is aggravated when yeast was grown at a low temperature, a condition that enhances proteotoxic stress. We observed that while ectopic expression of wild-type Rad23 improved the growth of the *rpn10Δrad23Δ* strain, expression of Rad23^UbLK0^ did not have any effect at 18°C or 30°C suggesting that the lysine residues in the UbL domain are required for Rad23’s function in proteasomal degradation (Fig. 5B). The importance of the lysine residues within the UbL domain for the ability of yeast to deal with proteotoxic stress was confirmed by the observation that expression of wild-type Rad23, but not Rad23^UbLK0^, restored growth of the *rad23Δ* strain in the presence of canavanine, an arginine analogue that causes severe proteotoxic stress (Fig. 5C). Also this phenotype was aggravated when canavanine sensitivity was analyzed in the *rad23Δrpn10-UIM* strain, which in addition to the Rad23 deletion also expressed an Rpn10 subunit that lacked the ubiquitin interaction motif (UIM). Thus, whereas wild-type Rad23 improved the growth of the *rad23Δrpn10-UIM* strain, the Rad23^UbLK0^ variant was unable to restore growth (Fig. 5D). This phenotype is similar to the one observed in wild-type yeast overexpressing Ubp12^FLAG^, which displayed reduced growth when yeast was cultured at low (18°C) or elevated (37°C) temperatures while a very mild effect was observed at 30°C suggesting that proteotoxic stress enhances the growth suppressive effect of Ubp12 overexpression (Fig. 5E). Together our data support the model that the ubiquitylation status of the UbL domain of Rad23 is important for Rad23’s function in proteasomal degradation while not being required for the non-proteolytic role of Rad23 in DNA repair. Thus, ubiquitylation of Rad23, which is regulated by Ubp12, stimulates proteasomal degradation of Rad23-dependent substrates and interference with this process results in stabilization of Rad23-dependent substrates, increased load of ubiquitylated cargo on Rad23 and an impaired ability to deal with proteotoxic stress.

## Discussion

In the present study, we show that Rad23 is ubiquitylated in its N-terminal UbL domain and that this modification is reversed by Ubp12, which we identified as a Cdc48-interacting DUB. We show that preventing ubiquitylation of Rad23 by either replacing acceptor lysine residues in the UbL domain or by overexpressing Ubp12 impairs the degradation of Rad23-dependent substrates. This defect in degradation of Rad23-dependent substrates is not caused by an impaired interaction with the proteasome but is accompanied by an increased load of ubiquitylated proteins that are bound to the UBA domains of Rad23. The presence of supraphysiological levels of ubiquitylated cargo on Rad23 that is proficient for binding to the proteasome suggests that Rad23 is unable to deliver ubiquitylated proteins at the proteasome.

It has been proposed that the presence of several ubiquitin shuttle factors with different, though partially overlapping, specificities might function as a regulatory layer that determines the targeting rate of distinct pools of substrate (Verma et al., 2004). Although there are some examples of posttranslational modifications and binding proteins that regulate the interaction of ubiquitin shuttle factors with the proteasome (Isasa et al., 2010, Liang et al., 2014, Sekiguchi et al., 2011), it is presently unclear if and how these processes are regulated by internal and/or external cues. It is feasible that, under certain conditions, Rad23 deubiquitylation behaves as a regulatory switch that stalls degradation of Rad23-dependent substrates thereby giving higher priority to substrates that reach the proteasome by different means, such as alternative shuttle factors. However, we consider such a role for ubiquitylation of Rad23 less likely as we would expect such a regulatory switch to operate before the interaction with the proteasome since Rad23 would otherwise still compete with other ubiquitin shuttle factors for binding to the proteasome and thereby also negatively affect degradation of their cargo. Our data, however, show that Rad23^UbLK0^ still interacts with the proteasome despite its impaired ability to facilitate degradation of substrates. Moreover, at any given time only a small fraction of Rad23 is modified with ubiquitin conjugates whereas interference with Rad23 ubiquitylation has a strong effect on degradation, which would be more consistent with a rather transient modification of Rad23 being required for degradation rather than a conditional regulatory switch.

Based on the present data, we propose an alternative, although not mutual exclusive, model in which a Cdc48-regulated deubiquitylation changes the properties of Rad23 in a dynamic fashion that contributes to the directionality in substrate delivery. Interestingly, ubiquitin is not only a signal for delivery of the proteins to the proteasome but also targets substrates to Cdc48 and other ubiquitin-binding proteins that act exclusively upstream of the proteasome (Schrader et al., 2009). Earlier work has shown that elongation of the ubiquitin chain ensures that substrates interact with Cdc48, Rad23 and the proteasome in a successive manner since the preferred ubiquitin chain length increases progressively along this route (Richly et al., 2005). However, while ubiquitin chain elongation may be important at the events that occur before the delivery at the proteasome, the proteasome itself has a broad specificity for ubiquitin modifications and hence additional cues may be required to facilitate the final delivery of the substrate (Grice and Nathan, 2016). Our data would be consistent with a model in which Rad23 deubiquitylation at Cdc48 switches Rad23 into a mode that allows delivery of its cargo to proteasomes. Although we have been unable to identify a specific ubiquitin ligase responsible for ubiquitylation of Rad23, it is feasible that Rad23 ubiquitylation is generally linked to association with the proteasome since a broad variety of ubiquitin ligases have been shown to interact with the proteasome (Xie and Varshavsky, 2000). It is also noteworthy that Rad23 can adopt open and closed conformations due to intramolecular interactions between its UbL and UBA domains and this process can be regulated by the ubiquitin receptor Rpn10 (Walters et al., 2003). It is tempting to speculate that the ubiquitylation status of the UbL domain modulates this process. Thus, our observation of the ubiquitylation status of Rad23 being governed by the Cdc48-associated Ubp12 and its implications for substrate delivery gives new insights in the molecular mechanism underlying the targeting of substrates for proteasomal degradation.

## Materials and methods

### Yeast strains and plasmids

Yeast strains were grown according to standard procedures on complete or synthetic media supplemented with 2% (w/v) glucose. Cycloheximide (Sigma) (100 μg/ml from a stock at 10 mg/ml in ethanol) or MG132 (Enzo) (50 μM from a stock at 10 mM in DMSO) was added when indicated. For the analysis of Rad23 ubiquitylation upon exclusive expression of Myc-Ubiquitin, the strain YD466, isogenic to SUB328, was used (Spence et al., 1995). Yeast strain SY980 (RPN10-UIM) was a gift from Suzanne Elsasser (Elsasser et al., 2004). A complete list of the yeast strains used in this study can be found in the supplementary information (**Suppl. Tab. 1**). The Ubp12-HA expression plasmid has been reported before (gift from Dr. Mafalda Escobar-Henriques) (Anton et al., 2013). The expression plasmid for Myc-tagged ubiquitin, the YEp112 plasmid (2μ, URA3, pCUP1-Ub) and plasmids for wild-type and lysine-less ubiquitin (Ub and UbK0; 2μ, LEU2) were provided by Drs. Jörgen Dohmen, Daniel Finley and Michael Glickman, respectively.

### Immunoprecipitations

Indicated strains were grown in 25 ml of appropriate medium and cells were harvested at an OD600 around 1, snap frozen stored if needed at −80 degree. Cells were harvested and resuspended in 500μl of ice cold lysis buffer (10mM Tris-Cl pH 7.5, 150mM NaCl, 0.5 mM EDTA, 0.5% NP-40, 20mM NEM, 10μM MG-132; for Ubp12 interaction: 0.05% Digitonin was used instead of NP-40) containing a protease inhibitor mix (complete, EDTA free, from Roche), and lysed with an equal volume of acid-washed glass beads (0.4–0.6 mm diameter) by vortexing five times for 1 min with 1 min intervals on ice. Lysates were clarified by centrifugation at 13000 rpm for 10 min at 4 °C. Protein concentrations were quantified by Bradford and lysates normalized to 1 mg/ml total protein. Ten μl lysate were removed as input sample. For immunoprecipitation, the following antibodies were used: Rad23 (goat polyclonal antibody, dilution 1:5000, R&D systems, A302-306), Dsk2 (rabbit polyclonal antibody, 1:5000, Abcam, ab4119-100), Cdc48 (rabbit polyclonal antibody, dilution 1:1000, Abcam, ab138298), and FLAG C-terminal (M2 mouse monoclonal antibody, 1:1000, Sigma, F3165). The indicated antibodies were added to 500 μl of lysates and incubated for 1 h on a rotating wheel at 4 °C. Then 50 μl Protein G Dynabeads (Invitrogen) washed three times in lysis buffer were added and incubated with the lysates for 1 hour on a rotating wheel at 4 °C. Using a magnet, beads were separated from the lysate and washed five times with 500 μl wash buffer. For anti-Flag IP, elution was performed with 1 μg/ml Flag peptide in wash buffer for 2 h at 4 °C. Otherwise, beads were incubated in 20 μl SDS buffer for 5 min at 95 °C. Input and eluate samples were analyzed by SDS-PAGE and Western blotting.

### Western blot analysis

The membranes were probed with antibodies directed against ubiquitin (rabbit polyclonal antibody, dilution 1:500, DAKO, Z0458), HA (mouse monocolonal antibody, dilution 1:5000, Biosite, 16B12), GFP (rabbit polyclonal antibody, dilution 1:5000, Molecular Probes, A6455), PGK1 (mouse monoclonal antibody, 1:5000, Life Technologies, H3420), FLAG N-terminal (mouse monoclonal antibody, M5, dilution 1:2000, Sigma, F4042), FLAG C-terminal (M2 mouse monoclonal antibody, 1:1000, Sigma, F3165), Cdc48 (rabbit polyclonal antibody, dilution 1:1000, Abcam, ab138298), Rad23 (rabbit polyclonal antibody, dilution 1:5000, gift from Dr. Kiran Madura), Rpt3 (rabbit polyclonal antibody, dilution 1:5000, Biomol, PW8250). Proteins were detected using either secondary anti-mouse, anti-goat or anti-rabbit IgG coupled to peroxidase (GE), chemiluminescent substrate (GE), and X-ray films or anti-mouse, anti-goat, or anti-rabbit IgG coupled to near-infrared fluorophores (LiCOR) and the Odyssey Infrared Imaging System. The latter system was also used for signal quantification.

### Cycloheximide chase assay

For cycloheximide chase analysis, exponentially growing cultures were adjusted to 100 μg/ml cycloheximide, and samples were taken at different time points. Protein extracts were prepared by lysis and precipitation in 27.5 % trichloroacetic acid and analyzed by SDS-PAGE and Wester blotting.

### Ubiquitylation assay

Yeast was transformed with an expression plasmid for His-tagged wild-type and lysine-less ubiquitin (gifts from Dr. M. Glickman and Dr. Daniel Finley). For precipitation of 6xHis-ubiquitin modified proteins under denaturing conditions a yeast culture of 200 OD600 units was harvested and lysed on ice in 6 ml lysis buffer (1.91 N NaOH, 7.5% (v/v) ß-mercaptoethanol). After lysis, 6 ml 55% trichloroacetic acid was added and the samples were vortexed and left for 15 min on ice. The samples were centrifuged at 13,000 rpm for 15 min at 4°C. The pellet was washed two times with 10 ml acetone after which the pellets were resuspended in 12 ml buffer A (6 M guanidiumchloride, 100 mM NaH_2_PO_4_, 10 mM Tris pH 8.0, 0.05% Tween 20). Imidazole was added to a final concentration of 20 mM together with 100 μl Ni-NTA magnetic agarose beads (Qiagen). The samples were incubated on a rotating wheel for 16 hrs at 4°C. The beads were washed two times while placed in a magnetic rack with buffer A and two times with buffer B (8 M urea, 100 mM NaH_2_PO_4_, 10 mM Tris pH 6.3, 0.05% Tween 20). The proteins were eluted by incubating the beads in 30 μl 1% sodium dodecyl sulfate for 10 min at 65°C. The eluates were dried in a speed-vac for 45 min, dissolved in 20 μl loading buffer and incubated for 10 min at 95°C.

### Growth assays

For spot assays six time serial dilutions of logarithmically growing cells were spotted on appropriate media containing 1.5 μg/ml canavanine, where indicated, and grown for 3 days. Growth at 18 °C was monitored after 4 days.

## Acknowledgments

The authors acknowledge the help of Tobias Schmidt and Tilman Kurz in early stages of the project, Drs. Jürgen Dohmen, Mafalda Escobar-Henriques, Suzanne Elsasser, Daniel Finley and Michael Glickman for reagents, and Dr. Florian Salomons for critical reading of the manuscript and technical input.

### Competing interests

No competing interests declared.

### Author contributions

Conceptualization: D.G., N.P.D.; Methodology: D.G., N.P.D.; Formal analysis and investigation: D.G., N.P.D.; Writing - original draft preparation: D.G.; Writing - review and editing: D.G., N.P.D.; Funding acquisition: D.G., N.P.D.; Supervision: N.P.D.

### Financial support

This work was supported by the Swedish Research Council (N.P.D.) and the Wenner-Gren Foundation (D.G.).

